# Environmental context, parameter sensitivity and structural sensitivity impact predictions of annual-plant coexistence

**DOI:** 10.1101/2023.02.13.528375

**Authors:** Alba Cervantes-Loreto, Abigail I. Pastore, Christopher R.P. Brown, Michelle L. Maraffini, Clement Aldebert, Margaret M. Mayfield, Daniel B. Stouffer

## Abstract

Predicting the outcome of interactions between species is central to our current understanding of diversity maintenance. However, we have limited information about the robustness of many model-based predictions of species coexistence. This limitation is partly because several sources of uncertainty are often ignored when making predictions. Here, we introduce a framework to simultaneously explore how different mathematical models, different environmental contexts, and parameter uncertainty impact the probability of predicting species coexistence. Using a set of pairwise competition experiments on annual plants, we provide direct evidence that subtle differences between models lead to contrasting predictions of both coexistence and competitive exclusion. We also show that the effects of environmental context-dependency and parameter uncertainty on predictions of species coexistence are not independent of the model used to describe population dynamics. Our work suggests that predictions of species coexistence and extrapolations thereof may be particularly vulnerable to these underappreciated founts of uncertainty.

## Introduction

The effects species have on one another are the result of multiple processes that often act simultaneously. In the case of competition between plants, examples include the depletion of local resources in the soil (Dybzinski & Tilman, 2007; Craine & Dybzinski, 2013), visits from shared pollinators (Lanuza *et al*., 2018), or the frequency and intensity of disturbance events (Pickett, 1980; Villarreal-Barajas & Martorell, 2009). Notwithstanding their importance, fully including all such phenomena in the study of plant dynamics is often impractical. Hence, it is more straightforward to treat these processes implicitly and model the relationship between interacting species phenomenologically, for example by fitting models that describe how the densities of intraspecific and interspecific neighbors change plant fitness and growth (Case, 1999; Connell, 1990; Goldberg, 1990; Adler *et al*., 2018; Hart *et al*., 2018).

Despite their “necessary incompleteness”, phenomenological models can accurately reproduce data observed in various natural systems and contexts (Hilborn & Mangel, 1997; Bolker, 2008; Houlahan *et al*., 2015). Perhaps more importantly, they are useful tools with which to make predictions that extend beyond the phenomena they describe (Broekman *et al*., 2019). Such predictions are possible because of the implicit assumption that models that reproduce the observed data faithfully also capture how the studied system operates (Marquet *et al*., 2015; Klir, 1985; Zeigler *et al*., 2000; Stouffer, 2022). For example, models that describe the effects neighboring plants have on each other can be used to make quantitative predictions about changes of biomass in the system (Godoy *et al*.,2020; Lai *et al*., 2022) or qualitative predictions such as whether or not co-occurring plant species can coexist (Levine & HilleRisLambers, 2009; Zepeda & Martorell, 2019).

The practicality of phenomenological models of plant competition, however, is a double-edged sword. Indeed, predictions made with them are subject to uncertainty arising from many distinct sources. One such source is environmental context dependency, or the extent to which the outcomes of species interactions change as a function of the abiotic conditions species experience (Bimler et *al.*, 2018; Chamberlain *et al*., 2014). Studies have found, for example, substantial evidence that interaction strengths between plants vary along environmental gradients (Bimler *et al*., 2018; Villarreal-Barajas & Martorell, 2009; Lanuza *et al*., 2018); interspecific interactions in particular can switch from competitive to facilitative when moving from favorable to harsh environments (Callaway *et al*., 2002; Maestre *et al*., 2005; Brooker *et al*., 2008; Maestre et al., 2009). This environment-driven variation can even lead to the identity of the competitive superior plant species changing depending on local abiotic conditions (Poorter & Lambers, 1986; Dybzinski & Tilman, 2007). Extrapolations from phenomenological models of plant competition can therefore be highly specific to the set of conditions under which models were parameterized (Bimler *et al*., 2018).

Model-based predictions are also subject to two forms of uncertainty that arise from the use of models themselves: parameter sensitivity and structural sensitivity. Parameter sensitivity refers to the sensitivity of model outputs to variation in parameter values, and exploring it constitutes a routine analysis in the domain of the biological sciences (Jørgensen & Bendoricchio, 2001; Terry et al., 2021). On the other hand, structural sensitivity characterizes how mathematical expressions that have similar phenomenological behavior can produce qualitatively different outcomes (Cordoleani *et al*., 2011; Myerscough *et al*., 1996; Aldebert & Stouffer, 2018). Parameter and structural sensitivity are often intertwined (Wood & Thomas, 1999), and both have been shown to drastically change model predictions in a vast array of biological systems (Cordoleani *et al*., 2011; Wood & Thomas, 1999; Poggiale *et al*., 2010; Fussmann & Blasius, 2005; Aldebert *et al*., 2016; Aldebert & Stouffer, 2018).

The interplay between environmental context dependency, parameter sensitivity, and structural sensitivity is rarely explored simultaneously, and to the best of our knowledge has never been explicitly explored for the case of models of annual-plant population dynamics. In this study, we therefore aim to understand how these three sources of uncertainty change predictions of a widely studied and vastly important ecological process: species coexistence. We focused our analysis on annual plants, which is a common natural system used to study species coexistence (Levine & HilleRisLambers, 2009; Godoy & Levine, 2014; Wainwright *et al*., 2019; Zepeda & Martorell, 2019). We assessed the empirical relevance of the three different sources of uncertainty by making coexistence predictions based around data from competition experiments between two annual-plant species conducted in two contrasting abiotic conditions. Our analyses provide evidence that uncertainty can radically change predictions made from a simple competition experiment, and highlights the importance of incorporating uncertainty from as many sources as possible when making model-based predictions.

## Methods

We will first provide a mathematical description of how to make and interpret coexistence predictions made with a population-dynamics model for two annual-plant species growing in proximity to each other. We then expand our framework to introduce alternative phenomenological models of density dependent seed production, and show how our framework can be used to make predictions using a different model for each species. Second, we describe how to use a Bayesian framework to parameterize the aforementioned phenomenological models to data from a set of competition experiments between two annual-plant species growing under two distinct abiotic conditions. Finally, we describe how we simultaneously explored how environmental context dependency, parameter sensitivity, and structural sensitivity impact predictions of species coexistence.

### Model-based predictions of species coexistence

We used the Cohen model (Cohen, 1966; Watkinson, 1980) to describe annual-plant population dynamics and as the starting point for our model-based predictions of species coexistence. This model predicts the density of seeds *N_i,t+1_* from species *i* in year *t* + 1 with:

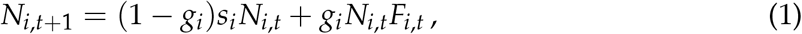

which is a function of (i) the density of seeds in the seed bank from the previous year (*N_i,t_*) that do not germinate yet remain viable bank in the seed bank (as weighted by *s_i_*, the annual rate of seed survival in the seed bank) and (ii) the density of seeds that germinate (determined by the germination rate *g_i_*) multiplied by the number of viable seeds produced per seed germinated, often called their “realized fecundity” (*F_i,t_*). The realized fecundity of species *i* can be accurately described by many different phenomenological forms (Law & Watkinson, 1987; Hart *et al*., 2018; Godwin *et al*., 2020; Stouffer, 2022). These phenomenological descriptions of *F_i,t_* generally try to capture the relationship between plant reproductive output and the densities of conspecific and heterospecific neighbors, but do not necessarily imply a hypothesis about the mechanisms underpinning this density dependence (Stouffer, 2022).

For example, *F_i,t_* can be given by the Beverton–Holt (a.k.a. reciprocal or inverse) model (Beverton & Holt, 1957), which in a two-species context equals:

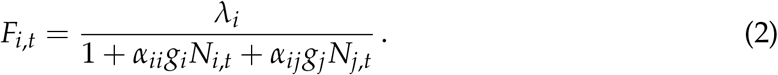

In this model, the per germinant fecundity of species *i* in the absence of density dependence is described by the parameter *λ_i_*, while the numbers of germinants of species *i* and *j* in year *t* are given by *g_i_N_i,t_* and *g_j_N_j,t_*, respectively. The density-dependent effects are captured by the interaction coefficients *α_ii_* and *α_ij_*, which describe the interaction strengths of conspecifics and heterospecifics, respectively. The Beverton–Holt model is a commonly used phenomenological model to make coexistence predictions and can be easily parameterized with empirical observations of annual plants growing in proximity to each other (Godoy & Levine, 2014; Godoy *et al*., 2014; Levine & HilleRisLambers, 2009; Hart *et al*.,2018).

#### Coexistence predictions

From the population dynamics that result from using Eqn. 1 and estimates of the relevant parameters of Eqns. 1 and 2, it is possible to predict if a pair of species can coexist. Multiple approaches exist to predict species coexistence (Chesson, 2000; 2018; Barabás et al., 2018; Saavedra *et al*., 2017; Letten *et al*., 2017). One of them is to directly evaluate, given the competitive constraints each species experiences, if the set of species’ intrinsic growth rates is feasible (i.e., if there exists an equilibrium point under which both species have positive densities; Rohr *et al*., 2014; Saavedra *et al*., 2017). To do so, it is necessary to derive the equations determining the equilibrium density for each species, which for species *i* is found at:

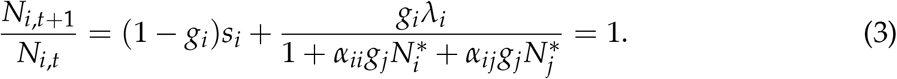

This equilibrium condition can be arranged to provide a linear equation in terms of densities:

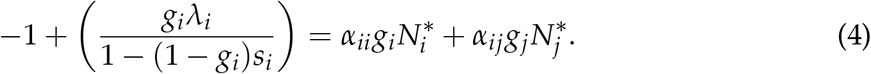

For simplicity, Eqn. 4 can be rewritten as:

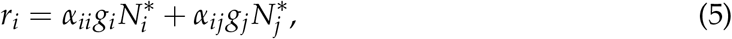

where *r_i_*, is the vital rate of species *i* that, at equilibrium, is compensated for by species interactions. Note that this vital rate *r_i_*, is a composite parameter that depends on the values of *s_i_*, *g_i_*, and *λ_i_*. Rather than call it an intrinsic growth rate, we purposefully refer to *r_i_* as a vital rate to emphasize that it does not correspond to the expected growth rate of species *i* when densities are vanishingly small nor when there are no interaction effects.

Equivalent expressions for species *j* can be derived from its equilibrium condition. The combined two-species equilibrium condition is:

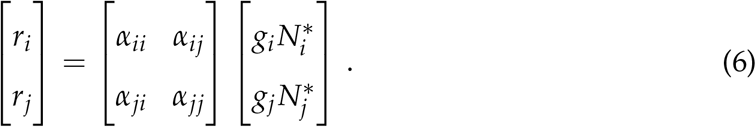

Given estimates of *r_i_*, *r_j_*, and the 2 × 2 matrix of interaction coefficients, predicted species densities at equilibrium can be determined by rearranging Eqn. 6 to:

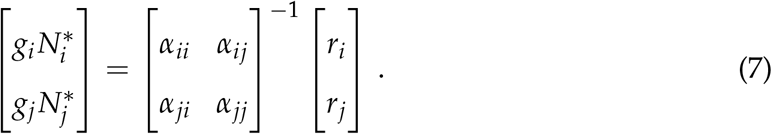

When predicted equilibrium densities for both species are positive, then the model-based prediction is that they can coexist (Rohr *et al*., 2014; Saavedra *et al*., 2017). In contrast, if one of the predicted equilibrium densities is less or equal to zero, then the model-based prediction is that one of the species will competitively exclude the other.

In practice, it is useful not only to determine if some particular values of *r_i_* and *r_j_* allow species to coexist, but to explore the full set of values of species’ vital rates that are compatible with species coexistence. This is often referred to as the *structural approach*, and is easily applicable to models of annual-plant population dynamics (Saavedra *et al*., 2017). The region of vital-rate parameter space where both species can have positive densities at equilibrium, given the constraints imposed through the interaction matrix, is called the feasibility domain (Rohr et al., 2014; Saavedra et al., 2017; Song et al., 2018; 2020). Biologically, a large feasibility domain means that competitive constraints are lax, and species can grow at different combinations of rates without excluding each other. In contrast, a small feasibility domain means that competitive constraints are harsh, and only a handful of vital rates allow their coexistence.

#### Biologically constrained feasibility domain

Importantly, locations in the vital-rate parameter space carry direct biological interpretations with them beyond whether or not equilibrium densities are feasible. Consider, for example, a vital-rate vector 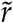 that allows for a vector of positive equilibrium densities *Ñ*\*. For any scalar value *x*, the proportional vector 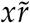 will also produce *xÑ*\* as a solution to Eqn. 7. However, it is reasonable to assume that there is an upper limit to species’ densities in nature (i.e., we do not expect species to achieve infinite local abundances). If a vital-rate vector leads to predicted densities beyond a particular species’ observable limit, we argue it may be mathematically feasible but should not be considered biologically feasible. The imposition of an abundance constraint such as this one will tend to create an upper bound on the vital rates that define the feasibility domain.

In addition, the Beverton–Holt model implicitly imposes further biological constraints on the values species’ vital rates can take. Recall that a species’ composite vital rate *r_i_* is a product of three biologically meaningful parameters. Those parameters have bounds themselves and when combined together they can further impact the values species’ composite vital rates can take. Specifically, *s_i_*, and *g_i_*, are proportions and can only have values between zero and one, while the per germinant fecundity in the absence of interaction effects *λ_i_* can only have positive values. By assuming density dependence for a given species follows the Beverton–Holt model, these parameter constraints together imply that vital rates *r_i_* < −1 are not biologically feasible. Indeed, all values of *r_i_* < −1 correspond to 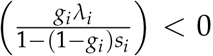, but for this inequality to be true we require either 1 – (1 – *g_i_*)*s_i_* < 0 or *g_i_λ_i_* < 0. Since *s_i_* and *g_i_* are proportions, 1 – (1 – *g_i_*)*s_i_* can never be lower than zero. Thus, the only way to obtain *r_i_* < −1 is for species *i* to have a negative intrinsic fecundity (*λ_i_* < 0), which is also not biologically plausible. In contrast, the Beverton–Holt model itself imposes no upper bound to species’ composite vital rates. Note that the lower bound of −1 and lack of an upper bound are specific to the Beverton–Holt model. As we will discuss later, different models of density-dependent fecundity will have different implicit model-based constraints on *r_i_*.

Building upon the feasibility domain defined within the structural approach, we define the biologically constrained feasibility domain as the region of parameter space where both species can have positive abundances given a) intraspecific and interspecific interactions, b) constraints on maximum species abundances, and c) vital-rate constraints imposed by each phenomenological fecundity model. In the two species case, the biologically constrained feasibility domain can also be thought of as an region that we denote with the symbol *β*. We estimated *A_β_*—the size of *β*—using Monte Carlo integration methods as described in Appendix S1, and show a visual example in Fig. 1.

**Figure 1:**
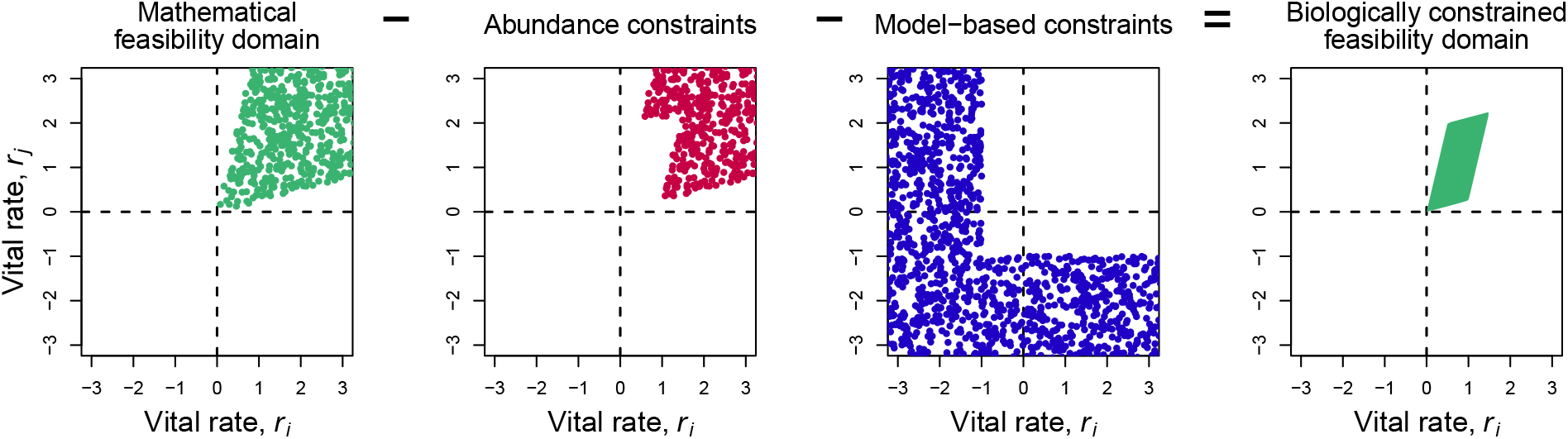
Estimation of the biologically constrained feasibility domain. A) We show the mathematical feasibility domain (green points) given a hypothetical competition matrix with intraspecific competition coefficients *α_ii_* = *α_jj_* = 1 and interspecific competition coefficients *α_ij_* = *α_ji_* = 0.5. Points sampled in this region of parameter space lead to both species having positive equilibrium densities. Note that, mathematically, this region extends infinitely in the positive quadrant. B) Some of these mathematically feasible points, however, may correspond to equilibrium abundances that are greater than empirically informed abundance constraints (red points). For this visual example, we restrict biologically sensible equilibria to have 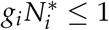 and 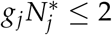. C) Part of the the parameter space may also fall outside model-based constraints (blue points); for example, when both species’ density dependence is described by the Beverton–Holt model *r_i_* ≥ −1 and *r_j_* ≥ −1. D) The space of the mathematical feasibility domain that does not overlap with any abundance or model-based constraints gives the biologically constrained feasibility domain (green area).

#### Relative coexistence ratio

As a basis of comparison, it is also useful to quantify the parameter space that allows both species to grow in monoculture (i.e., when they have no niche overlap). Importantly, this parameter space can also be expressed as an area, and for equivalent reasons to those described above this area is also subject to both abundance and model-based constraints. We denote this region with the symbol *γ*, and mathematical details of how we calculate its size, *A_γ_* are described in Appendix S2. By comparing the size of the parameter space where both species can feasibly coexist (*A_β_*) to the size of the space where species can grow in monoculture (*A_γ_*), we can quantify the importance of interspecific interactions relative to intraspecific interactions in determining the vital rates consistent with biologically plausible coexistence. This comparison can be expressed as a ratio *ρ* = *A_β_*/*A_γ_* that we call the relative coexistence ratio. When the ratio *ρ =* 1, then species coexistence is as achievable as species growing in monoculture; when the ratio *ρ* < 1, then the parameter space where the two species can coexist is smaller than the parameter space where each species can grow in monoculture, and it is harder for them to coexist because of their interspecific interactions; finally, ratios *ρ* > 1 imply that species facilitate each other, and it is easier for them to coexist because of their interspecific interactions.

#### An alternative model of density dependence

The Beverton–Holt model is only one of many phenomenological models used to describe density-dependent performance of annual plants. There is no hard-and-fast rule for how to choose the appropriate phenomenological model to describe the effect of species interactions, and it is often a choice governed by mathematical convenience (Stouffer, 2022), the type of study system (Godwin et al., 2020), and the governing paradigm around species interactions (Martyn *et al*., 2021). One alternative to the Beverton–Holt model is the Ricker model:

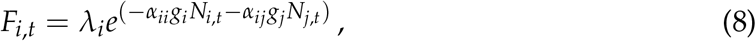

where the interpretation of the parameters remains the same as previously described (Ricker, 1954a;b). The Ricker model is known to be a biologically plausible and versatile model to quantify density dependence in annual-plant communities, plus it has the virtue of being better than the Beverton–Holt model at capturing both competitive and facilitative interactions (Mayfield & Stouffer, 2017; Bimler et al., 2018; Martyn et al., 2021; Stouffer, 2022). When discrete response variables are modeled as Poisson or negative-binomial random variates, for example within a generalized linear models, the default log link function also implicitly imposes a Ricker function on the model being fit (Rao et al., 2010; O’Hara & Kotze, 2010; Mayfield & Stouffer, 2017; Stouffer et al., 2018).

Similar to the process we followed from Eqns. 4 to 7, the Ricker model has its own composite vital rate at equilibrium. For species *i*, this is given by:

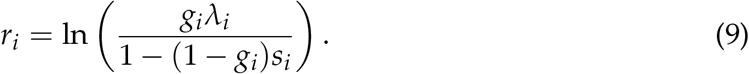

This expression necessarily implies that the model-based constraints on vital rates are fundamentally different when using the Ricker model than when using the Beverton–Holt model: it has no model-based constraints on the lower and upper bounds of species vital rates.

#### Multi-model predictions of species coexistence

By parameterizing a phenomenological model of plant competition, like the Beverton–Holt (Eqn. 2) or Ricker models (Eqn. 8), we get a description of how seed set varies with neighbor density. Perhaps more importantly, by linking these models to Eqn. 1 we can make predictions about whether or not two species are able to coexist. So far, we have been working with the implicit assumption that the same model describes the density dependence of the two species. However, given the lack of a clear mechanistic underpinning with which to justify using one model over another (Stouffer, 2022), there is no *a priori* reason to assume this is the case. This is particularly true when the models are parameterized with experimental data and from a statistical perspective appear to provide comparable fits (Hart *et al*., 2018).

To predict coexistence using a different density-dependence model for each species, we still must solve for species equilibrium abundances as a function of vital rates and competition (Eqn. 7). Unlike the single-model approach, each species’ vital rate has its own formulation based on the model used to describe each species’ realized fecundity (Appendix S1: Table S1). As in the previous approach, if Eqn. 7 predicts positive abundances for both species within the bounds of each species’ model-based and abundance constraints, species are predicted to coexist.

### The interplay of different forms of uncertainty

#### Data

To assess the relevance of different sources of uncertainty in an explicit empirical context, we made coexistence predictions using parameter estimates inferred directly from a set of competition experiments. During 2017, we conducted pairwise competition experiments between two annual-plant species, *Goodenia rosea* and *Trachymene cyanopetala*. These experiments took place in the West Perenjori Nature Reserve in Western Australia (−29.479°S, 116.199°E). The reserve is dominated by York gum-jam woodlands, which support an understory of mixed native and exotic annual grasses and forbs (Dwyer et al., 2015).

Using locally collected seeds, we conducted competition experiments using a response–surface experimental design which has the advantage of being able to accurately distinguish intra and interspecific competition (Inouye, 2001; Hart et al., 2018). Within this design, we varied the densities of both species independently by using treatments that combined the two species at two or more densities. To also study how abiotic conditions changed coexistence predictions, the experiments took place in two contrasting environments: an *open* environment, where plants interacted at least 1 m away from *woody* debris and a woody environment, where plants interacted within 30 cm of woody debris.

To implement our response–surface experiments, in October 2017 we first weeded out all aboveground biomass of plants inside of circular plots with a 7.5 cm radius. This neighborhood radius has been shown to be sufficiently large to capture local plant–plant interactions within the study system (Martyn *et al*., 2021). Each plot was then sown at one of four different densities of each species as a focal: an invasion density where only five seeds were sown (thinned to one individual after germination), low density (15 seeds in total), medium density (30 seeds in total), and high density (60 seeds in total). We varied the densities with which the non-focal species was sown across three treatments: absent (0 seeds in total), medium (30 seeds in total), or high (60 seeds in total). Treatments therefore consisted of combinations of the density of each species sown as a focal, the density of each species sown as a neighbor, and the environment where interactions took place. We had eight replicates per treatment, which yielded 256 total plots. Seeds were allowed to germinate, and we thinned plots in composition in July 2018 (i.e., we weeded out neighboring plants from species that were not sown) and counted the abundance of neighbors removed during weeding. We then collected the seeds produced after the growing season in October 2018 and also counted the number of conspecific plant individuals and heterospecific plant individuals in the plot at the time of seed collection.

Finally, we relied on a different set of data to obtain estimates of the survival and germination rates for both of our focal species in the field and the maximum abundance each species could achieve in the neighborhood radius where interactions took place (Towers *et al*., 2022). We show these values of seed survival rate, germination rate, and maximum abundance per species in Appendix S3.

#### Statistical inference

We fit Eqns. 2 and 8 separately for both of our focal species in order to get the relevant parameter estimates necessary to make coexistence predictions. For both species, we fit these non-linear models with a Bayesian framework using Hamiltonian Monte Carlo methods. We used Bayesian inference to explicitly incorporate the uncertainty surrounding model parameters in probability distributions (McElreath, 2018). Across all models, we explicitly accounted for the environment where seeds were sown in our parameter estimates. For all of the parameters across all models, we did this by treating the *woody* treatment as a dummy variable W that takes the value *W* = 0 in the *open* conditions and *W* = 1 in the *woody* conditions.

For all models and all environmental conditions, we constrained the fecundity in the absence of density dependence to be positive in order to keep our predictions biologically plausible. Across all model fits, we included an extra interaction coefficient (*α_ik_*) to account for the effects of non-focal plant species that were weeded out in the experiment. We fit this extra interaction term to improve the parameter estimates related to our focal species, but because we do not know those species’ other parameter values we could not model their corresponding coexistence outcomes. To ease later interpretation, we constrained all focal–focal interaction coefficients to be positive (i.e., competitive). This only led to minor variation in inferred parameter values as only the posterior distributions of the “other” interaction coefficients showed a strong tendency toward values ≤ 0 (Appendix S4: Figs. S1 & S2).

Since seeds produced per focal individual is a discrete count response variable, we modelled these outcomes as negative-binomial variates. We used a negative binomial instead of a Poisson model because the former provided significantly improved model fit, indicating a degree of over-dispersion in the empirical data. To ensure consistency across our different linear and non-linear models, we fit all models using the negative binomial family and an identity link function.

We fit all models using the function *brm* from the package *brms* (Bürkner, 2017) in the statistical program R version 4.0.2 (R Core Team, 2013). As non-linear models, *brms* required explicit specification of the prior distributions for all parameters except the dispersion parameter in the negative-binomial distribution (for which we used the default prior); across all models, we therefore used wide priors for intrinsic fecundities and weakly informative priors for the interaction coefficients. As a representative example, the full Bayesian description of the Beverton–Holt model for species i is:

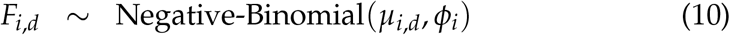

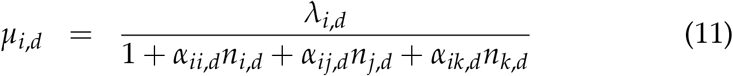

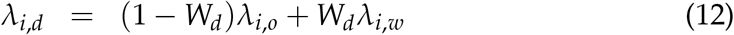

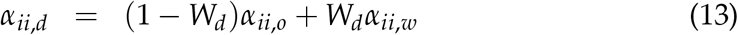

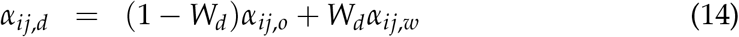

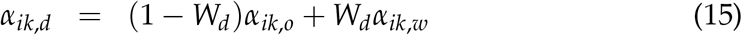

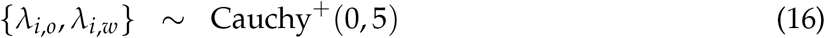

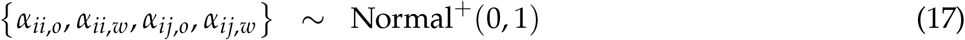

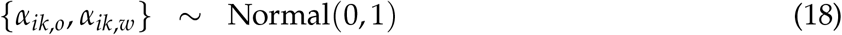

where the subscript *d* indicates each focal observation, the subscript o indicates parameters applicable in the *open* environment, the subscript *w* indicates parameters applicable in the woody environment, and *n_i,d_* is the count of species *i* in the plot corresponding to focal observation *d*. Only Eqn. 11 changes when moving from the Beverton–Holt to the Ricker model.

For each model, we ran four chains with a warmup of 2000 iterations and 2000 sampling iterations. We determined convergence when trace plots were well mixed and stationary and when the Gelman–Rubin convergence diagnostic (Rhat) was less than 1.05 for all parameters (Vehtari et al., 2021). We compared the fits of the models for each focal species using the Leave-One-Out cross-validation Information Criteria (LOOIC). This goodness of fit measure is used for estimating the out-of-sample prediction accuracy of Bayesian models and provides a measure of model fit that is penalized for the number of model parameters. As with other information criteria, lower values of LOOIC correspond to better supported models. Additionally, LOOIC is more robust for models with weak priors or influential observations compared to other information criteria (Vehtari *et al*., 2017). We calculated out-of-sample deviance separately for models in *open* and *woody* environments because LOOIC is calculated additively over observations. We also calculated Akaike weights for each model in each environmental condition, which can be interpreted as an estimate of the probability that the model will make the best predictions of new data, based on the set of models considered (McElreath, 2018). As one final basis of comparison, we also fit a “null” model where all interaction coefficients were fixed to zero as a demonstration that the models were capturing meaningful signals of density dependence in the empirical data.

#### Predictions incorporating uncertainty

To study how model formulation changes predictions of species coexistence, we used our framework to make predictions using median parameter estimates and a different model per species (Eqn. 2 or 8). We examined all the possible combinations of each species’ density dependence being defined by a different model, which yielded a total of four predictions. Furthermore, we also explored how abiotic conditions change predicted co-existence outcomes by making predictions using median parameter estimates in the *open* and *woody* conditions. We therefore had a total of eight different scenarios from which to make predictions (four for each environmental condition), as well as the corresponding size of *β*, size of *γ*, and value of *ρ*.

To incorporate parameter uncertainty, we also made predictions using 4000 draws from the parameters’ posterior distributions. For each of the eight predictions made using median parameter values, this gives us 4000 additional coexistence predictions. Posterior distributions of parameters contain the relative plausibility of different parameter values, conditional on the data and the model used (McElreath, 2018). This approach yielded a posterior distribution of coexistence outcomes, as well as distributions of the size of *β*, the size of *γ*, and the value of *ρ*.

For each model combination and each environmental condition, we lastly determined the proportion of posterior draws that predicted coexistence and competitive exclusion driven by *G. rosea* or *T. cyanopetala*. When the predicted outcome was competitive exclusion, this implies that the empirical expectation is a monoculture of the “dominant” species. Therefore, we also qualified these single-species dominant equilibria with whether or not the species was expected to be present at densities below or above the abundance-based constraints we considered elsewhere.

## Results

### Model fits

Model comparison using LOOIC showed that the Ricker model was the preferred model overall for density dependence of *G. rosea* and the Beverton–Holt model was the preferred model overall for *T. cyanopetala* since they consistently had the lowest LOOIC score (Table 1). However, Akaike weights showed that there was support for both models across the *open* and *woody* environmental conditions. Likely owing to the particularly weak interactions experienced by *T. cyanopetala* in the *woody* environment (Appendix S4: Figs. S1 & S2), the null model attained non-trivial model weight; however, the null model received limited Akaike weight across all the data, supporting the notion that the Ricker and Beverton–Holt models are indeed capturing meaningful signals of density-dependent fecundity for both two focal species.

**Table 1:** Model comparison for data collected in the *open* environment, the *woody* environment, and for all data together. LOOIC (leave-one-out cross-validation information criteria) penalizes models for the number of parameters, and the lowest value reflects the best-performing model. The resulting Akaike weight for each model is an estimate of the probability that the model will make the best predictions of new data across the the set of models considered.

| Species | Model | Open |  | Woody |  | All |  |
| --- | --- | --- | --- | --- | --- | --- | --- |
|  |  | LOOIC | Weight | LOOIC | Weight | LOOIC | Weight |
| *G. rosea* | Null | 691.96 | 0.01 | 693.45 | 0.00 | 692.70 | 0.00 |
| *G. rosea* | Beverton–Holt | 682.73 | 0.55 | 682.95 | 0.39 | 682.84 | 0.47 |
| *G. rosea* | Ricker | 683.18 | 0.44 | 682.10 | 0.60 | 682.64 | 0.52 |
| *T. cyanopetala* | Null | 946.47 | 0.00 | 943.19 | 0.06 | 944.83 | 0.01 |
| *T. cyanopetala* | Beverton–Holt | 934.95 | 0.66 | 937.60 | 0.92 | 936.28 | 0.88 |
| *T. cyanopetala* | Ricker | 936.33 | 0.33 | 944.52 | 0.03 | 940.42 | 0.11 |

### Structural sensitivity & environmental context dependency

As a starting point, we focus on predictions made using median parameter values but while varying the models used for density dependence and the environment in which the species were competing. Here, we found that inferences related to the biologically constrained feasibility domain, and how it varied across environments, were clearly contingent on the model formulation used for both specfies. The location of the biologically constrained feasibility domain showed clear shifts when moving from the *open* to *woody* environment (Fig. 2). Effectively all model combinations indicated that interactions allowed for greater variation in the vital rate of *G. rosea* in the open environment and for greater variation in the vital rate of *T. cyanopetala* in the *woody* environment while still maintaining coexistence. Three out of four model combinations also gave strong indications that the size of the biologically constrained feasibility domain was larger in the woody environment than in the *open* environment (Fig. 3). Similarly, all model combinations indicated that the size of the biologically constrained feasibility domain was (i) larger than the size of the area in monoculture in the *open* environment but (ii) smaller than the size of the area in monoculture in the *woody* environment (Fig. 4).

**Figure 2:**
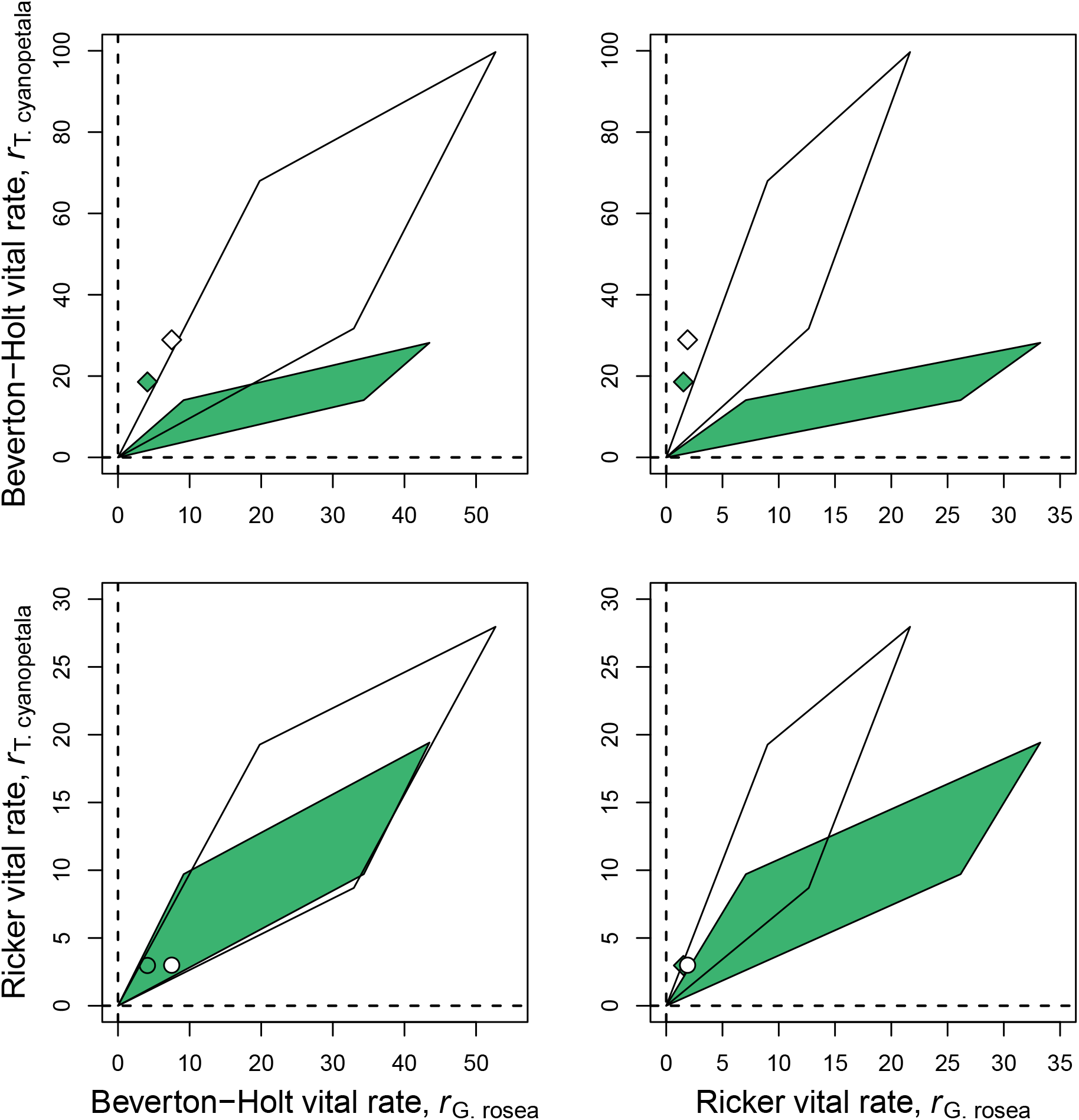
Biologically constrained feasibility domains and species’ inferred vital rates in the *open* and *woody* environments for different model combinations. For each combination of models used to quantify species’ density-dependent fecundity, we show the inferred feasibility domain and species vital rates using the median parameter values inferred from the experimental data. In each panel, the white area is the biologically constrained feasibility domain in the *open* environment and the green shaded area is the same in the *woody* environment. The white point corresponds to the inferred vital rates in the *open* environment and the green point the same in the *woody* environment. Points that are circles imply predicted coexistence (i.e., the vital rates fall inside the biologically constrained feasibility domain); points that are diamonds correspond to a prediction of competitive exclusion (i.e., the vital rates fall outside the biologically constrained feasibility domain).

**Figure 3:**
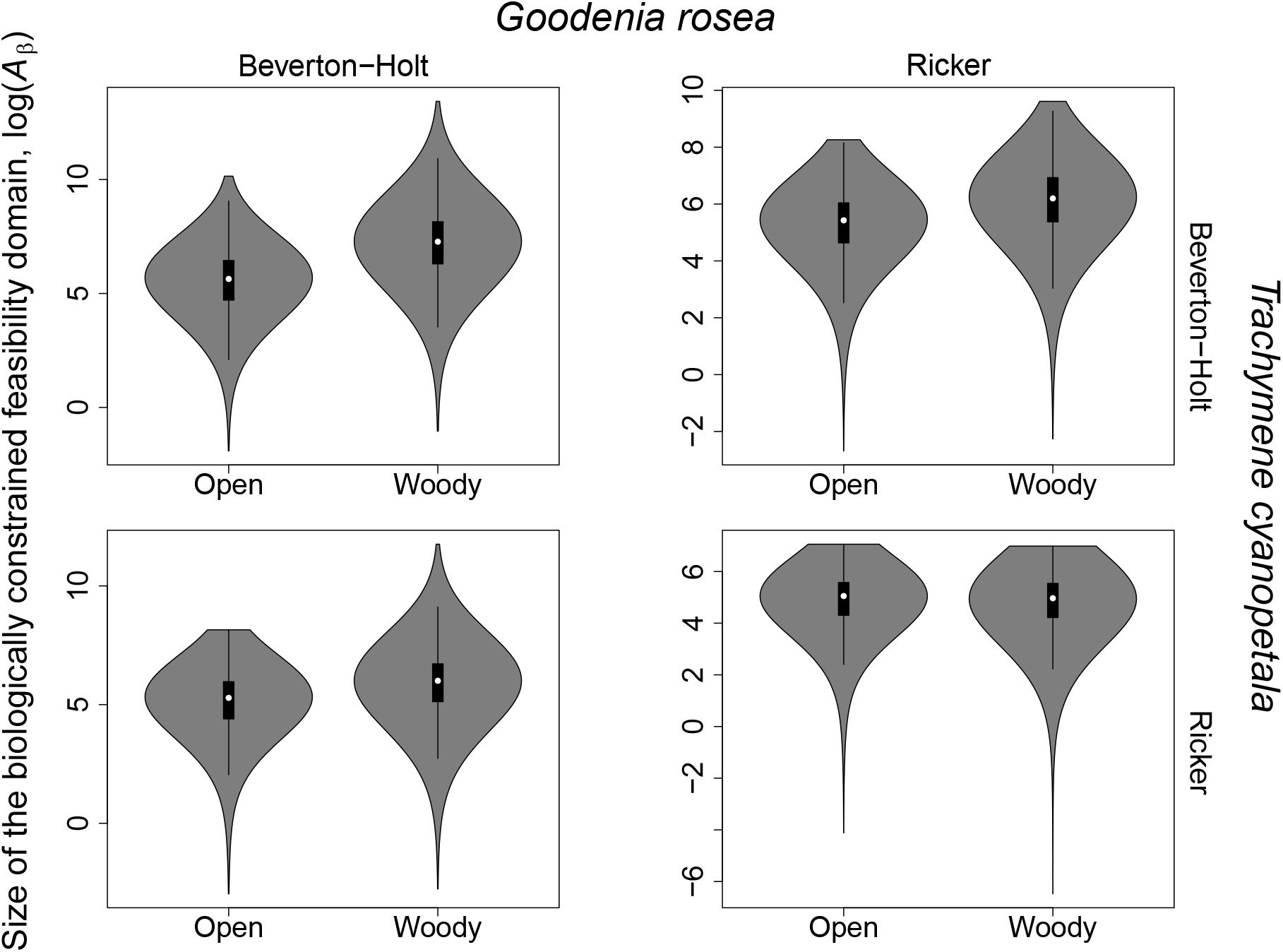
Estimated size of the biologically constrained feasibility domain (*A_β_*) in the *open* and *woody* environments for different model combinations. Each column corresponds to a different model used to capture density-dependent fecundity of *G. rosea* whereas each row corresponds to a different model used to capture density-dependent fecundity of *T. cyanopetala*, as indicated on top and at right. Within each panel, we show box-and-whisker plots for *A_β_* estimated from each draw from parameters’ posterior distributions. In all cases, the box covers the 25th-75th percentiles, the middle line marks the median, and the maximum length of the whiskers is 1.5 times the interquartile range. Underneath the box-and-whisker plots, we show violin plots which demonstrate the full posterior distributions of the quantities in question. Due to their large underlying variation, we logarithmically transform the values of *A_β_* prior to plotting.

**Figure 4:**
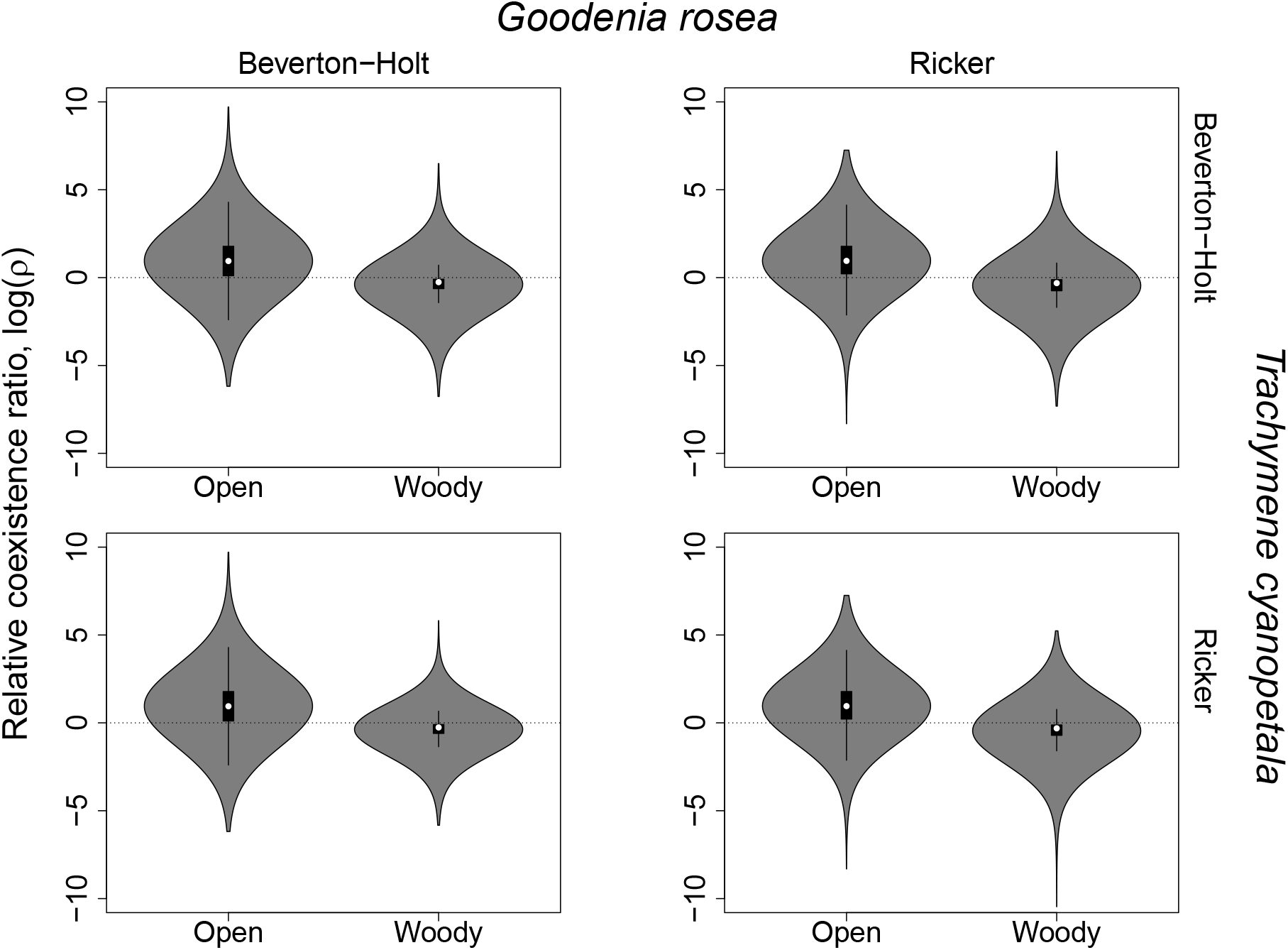
Estimated relative coexistence ratio *ρ* in the *open* and *woody* environments for different model combinations. Each column corresponds to a different model used to capture density-dependent fecundity of *G. rosea* whereas each row corresponds to a different model used to capture density-dependent fecundity of *T. cyanopetala*, as indicated on top and at right. Within each panel, we show box-and-whisker plots for sizes estimated from each draw from parameters’ posterior distributions. In all cases, the box covers the 25th-75th percentiles, the middle line marks the median, and the maximum length of the whiskers is 1.5 times the interquartile range. Underneath the box-and-whisker plots, we show violin plots which demonstrate the full posterior distributions of the quantities in question. Due to their large underlying variation, we logarithmically transform the values of *ρ* prior to plotting. The dotted line at log(*ρ*) = 0 therefore indicates when the two domains have equal sizes.

Predictions of species coexistence using median parameter estimates were also contingent on the model formulation used for both species in both environments (Fig. 5). In the *open* environment, the model used for *G. rosea* did not impact these predictions; however, we predicted *T. cyanopetala* would outcompete *G. rosea* when using the Beverton–Holt model for *T. cyanopetala* and predicted coexistence when using the Ricker model for *T. cyanopetala*. In the *woody* environment, the prediction using median parameter values was that *G. rosea* would outcompete *T. cyanopetala* as long as the Ricker model was used for *G. rosea* or when both species were modelled with the Beverton–Holt model; when the Beverton-Holt model was used for *G. rosea* but the Ricker model was used for *T. cyanopetala*, the prediction using median parameter values was that the two species would coexist.

**Figure 5:**
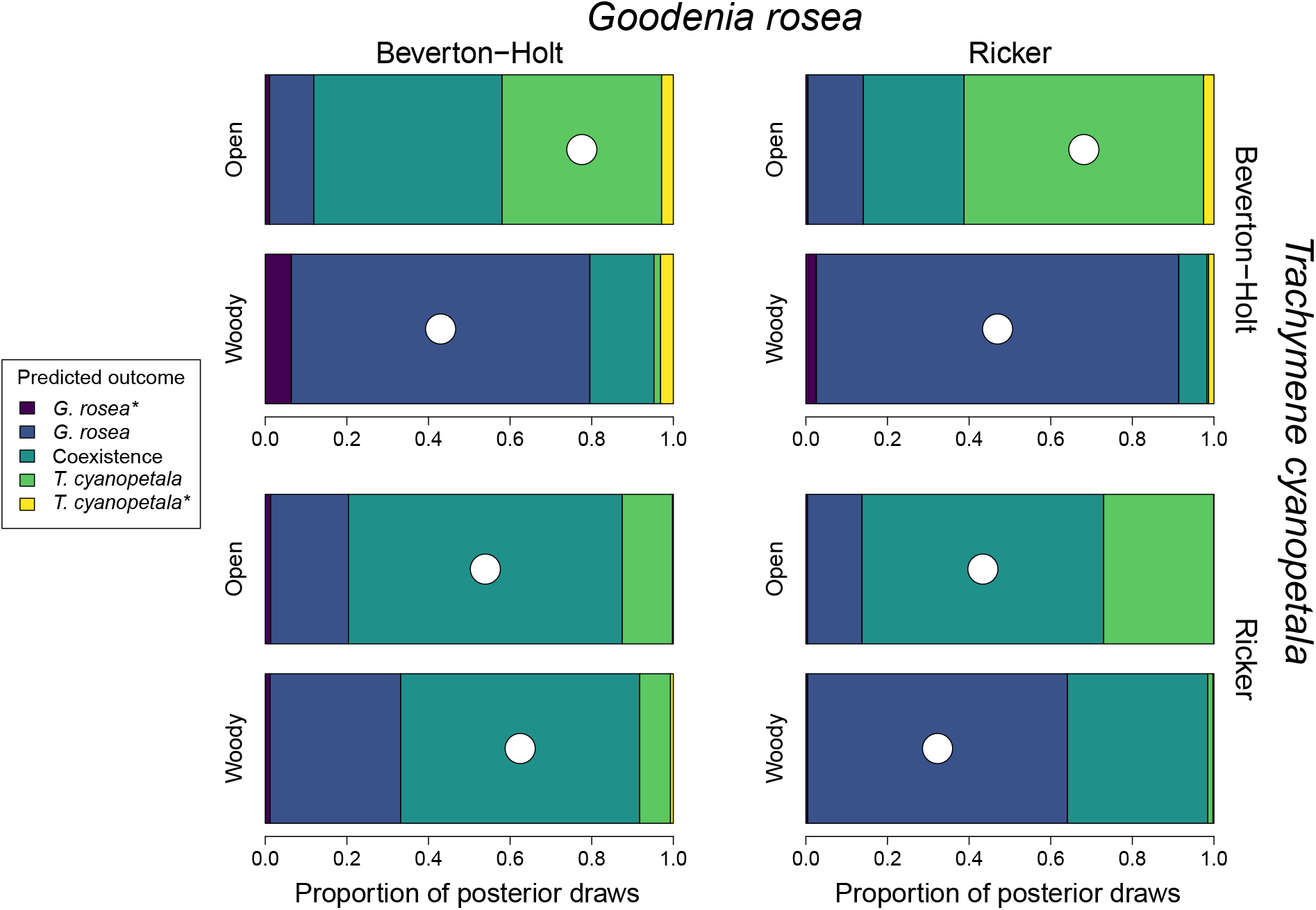
Predicted outcome of competition between the two focal species in the *open* and *woody* environments for different model combinations. Each paired set of horizontal bars corresponds to a different model combination; labels on the top indicate the model used for the species *G. rosea*, and labels on the right show the model used for *T. cyanopetala*. Horizontal bars within those panels show the proportion of posterior draws that predict: (i) competitive dominance of *G. rosea*, coexistence, or competitive dominance of *T. cyanopetala* as indicated with the color bar in the legend to the left. Outcomes highlighted with * indicate predictions of competitive dominance but at higher than realistic abundances. In each bar, we also highlight with a white circle the prediction corresponding to the median parameter values across all posterior draws.

### Structural sensitivity, environmental context dependency, & parameter sensitivity

Predictions made with the full posterior distributions of parameter values could be quantitatively and qualitatively very different than predictions made with median parameter values (Figs. 3, 4, & 5). Some model combinations appeared to induce greater variation in the location of the biologically constrained feasibility domain whereas other model combinations appeared to induce greater variation in the species’ inferred vital rates (Appendix S5: Figs. S1 & S2). The extent of parameter sensitivity depended on the models used and the environment where interactions took place. Importantly, we also noticed that the prediction made using median parameter values did not always correspond to the outcome that was predicted to be most likely across the full posterior distribution of parameter values (Fig. 5). There was a slightly higher tendency for model combinations including the Beverton–Holt model to predict mono-species dominance at a density greater than that given by our abundance-based constraints; however, these predictions were rare overall.

When the two species interact in the *open* environment, the most common prediction across posterior draws was that the two plant species would coexist (49% of all posterior draws). That said, the proportion of posterior draws that predicted coexistence ranged from 25% to 67% depending on the models used to capture density dependence. In the open environment and across all model combinations, 34% (range: 12–58%) of posterior draws predicted that *T. cyanopetala* would competitively exclude *G. rosea* whereas 14% (range: 11–19%) predicted the reverse. The *open* environment therefore was the environment in which *T. cyanopetala* appeared to have a competitive advantage.

When the two species interact in the woody environment, the most common prediction across posterior draws and model combinations was that *G. rosea* would competitively exclude *T. cyanopetala* (mean: 64%; range 32–88%). The next most likely predicted outcome was coexistence (mean: 29%) though this varied substantially across model combinations, from a low of 7% to a high of 59%. Competitive exclusion of *G. rosea* by *T. cyanopetala* was predicted to be rare across posterior draws (mean: 3%); however, it was predicted to occur for >7% of posterior draws when the Beverton–Holt model was used for *G. rosea* and the Ricker model was used for *T. cyanopetala*. The *woody* environment therefore was the environment in which *G. rosea* appeared to have a competitive advantage.

## Discussion

Our results show that using phenomenological models to predict whether or not a pair of species can coexist is far from trivial. Indeed, even seemingly subtle differences led to predictions of both coexistence and competitive exclusion based around the exact same experimental data. Even in the cases where we predicted the most likely outcome in a given environmental condition was competitive exclusion, which of the two species was predicted to be dominant could vary depending on the models being used.

Given our experimental data, both the Beverton–Holt and Ricker models received statistical support based on how they captured variation of seed sets with increasing neighbor density (Table 1). It is often the case in annual-plant studies that more than one phenomenological model has statistical support for different species or sites (Levine & HilleRisLambers, 2009; Mayfield & Stouffer, 2017; Bimler *et al*., 2018; Martyn *et al*., 2021). Nonetheless, exploring predictions made by more than one model is not common practice in the study of species coexistence, unlike the study of other ecological processes like predator–prey dynamics (Myerscough *et al*., 1996; Fussmann & Blasius, 2005; Aldebert & Stouffer, 2018). By limiting our predictions to a single type of phenomenological model, we not only have to ignore other models that share statistical support, but we also limit our understanding of how model formulation itself changes our predictions.

### Small differences, amplified

We found clear indications that that predictions of species coexistence using the annual-plant model are structurally sensitive. In both environments, predicted coexistence varied considerably depending on the models used to quantify density dependence (Fig. 5). Our study thus provides another clear example where predictions made with different models can be quantitatively similar (e.g., in terms of their fit to data) and still have different qualitative behaviors, one of the hallmarks of structural sensitivity (Cordoleani *et al*., 2011). Structural sensitivity can arise because slight perturbations in model formulation become largely amplified (Cordoleani *et al*., 2011; Wood & Thomas, 1999), and predictions can be structurally sensitive in different ways depending on the qualitative behavior examined (Cordoleani *et al*., 2011; Aldebert & Stouffer, 2018). In the case of coexistence outcomes, predictions require the choice of a model to use for each species, which can further amplify small differences between models.

In the *open* environment, we predicted coexistence was the most likely outcome in three out of the four model combinations we explored (Fig. 5). However, this depended most strongly on the model used to model density-dependence of *T. cyanopetala:* when using the Beverton–Holt model, competitive dominance of *G. rosea* by *T. cyanopetala* was predicted to be more likely. Overall, both the Beverton–Holt and the Ricker models predicted intraspecific interactions to be larger than interspecific interactions for both species in the *open* environment (Appendix S4: Figs. S1 & S2), which tends to promote coexistence since species limit themselves more than they limit others (Chesson, 2000; Adler *et al*., 2018). Furthermore, the intrinsic fecundity of *T. cyanopetala* was predicted to be larger given the Beverton–Holt model than given the Ricker model (Appendix S4: Fig. S2). Thus small differences between models resulted in a more restricted feasibility domain, and a shift in the position of species vital rates in parameter space, compared to other model combinations. Summed together, these changes caused the vital rates to fall outside of the biologically constrained feasibility domain and ultimately to a prediction of competitive exclusion of *G. rosea* by *T. cyanopetala* (Fig. 5).

### The importance of environmental context

The extent of structural sensitivity observed in our focal system changed in the *woody* environment. When we made predictions using median parameter estimates in the *woody* environment, three out of four model combinations predicted competitive exclusion of *T. cyanopetala* by *G. rosea* (Fig. 5). Intriguingly, this occurred despite the fact that the size of the biologically constrained feasibility domain in the *woody* environment appeared to be larger than its size in the *open* environment (Fig. 3). The increased likelihood of predicting competitive exclusion compared to the *open* environment seemed to be driven by several factors. One of them is that inferred interspecific interactions were stronger than intraspecific interactions for *G. rosea* when using both models (Appendix S4: Fig. S1), which resulted in a smaller *β* in the *woody* environment. However, since intraspecific interactions were generally inferred to be weaker in the *woody* environment (Appendix S4: Figs. S1 & S2), the space of vital rates allowing feasible coexistence compared to the space allowing monocultures decreased (Fig. 4).

Furthermore, both species in the *woody* environment were predicted to have lower fecundities in the absence of competition compared to the *open* environment, which supports observations from previous empirical studies in this system that observed lower plant abundances in this system when growing under litter (Wainwright *et al*., 2017b). Of the two species, G. rosea was predicted to experience a sharper reduction in seed set by both models (Appendix S4: Fig. S2). Even though the presence of woody debris in semiarid systems can reduce solar radiation and ameliorate drought stress (Wainwright *et al*., 2017b), our results suggest that these two species are less likely to coexist in woody environments due more to an environmentally induced shift in interaction strengths as opposed to a concerted change in the two species’ fitness differences.

Our results provide another example where interaction strengths, and thus coexistence predictions, change with environmental conditions, a result that has been empirically demonstrated before in this system (Wainwright *et al*., 2017a; Bimler *et al*., 2018) and others (Matías *et al*., 2018; Van Dyke *et al*., 2022). Other studies have documented that the fecundities of both focal species indeed change while growing inside coarse woody debris, but spatial mechanisms of coexistence have not been found (Towers *et al*., 2020). Importantly, the overall extent of environmental context dependency in our experimental system is influenced by the models used to quantify density dependence for both species. That is, the apparent effect of abiotic conditions can be enhanced or diminished in predictions of species coexistence due simply to one’s choice of phenomenological model(s).

### Parameter uncertainty and the probability of predicting species coexistence

Using a Bayesian approach to fit models to data also allowed us to have a better understanding of the parameter uncertainty associated with our predictions. Our results showed that estimating pairwise coexistence only using median estimates of parameter values might overlook instances where the uncertainty encompasses outcomes different to the median prediction (Fig. 5). Previous studies have also incorporated parameter uncertainty in coexistence predictions by propagating standard errors (Matías *et al*., 2018) or bootstrapping observations (García-Callejas *et al*., 2020). However, these approaches were only incorporated to show the robustness of predictions rather than to examine the causes and effects of parameter uncertainty in predictions of species coexistence.

Our results also show that even when we predicted competitive exclusion, the species we predicted to be competitively excluded varied across posterior draws (Fig. 5). Other studies that have incorporated posterior distributions of parameter values in coexistence predictions have also encountered this uncertainty regarding the outcome of competition (Terry *et al*., 2021). However, they also found that posterior predictions mostly agree with predictions using median parameter values (i.e., species were confidently coexisting or not; Terry *et al*., 2021). Our results did not show as clear differences, particularly in the *open* environment where species inferred vital rates were particularly close to the coexistence boundary (Appendix S5: Fig. S1).

Importantly, the effect of parameter sensitivity on predictions of species coexistence has been mostly interpreted as statements of uncertainty in the underlying data, rather than implying a probabilistic outcome (Terry *et al*., 2021; Matías *et al*., 2018). Our study goes beyond that interpretation by combining model weights and posterior predictions to calculate the probability of predicting coexistence given the phenomenological models used to quantify density dependence. Given the uncertainty associated with our predictions, our results suggest that coexistence between *G. rosea* and *T. cyanopetala* is at least plausible in the *open* environment while being virtually impossible in the *woody* environment (Fig. 5).

## Conclusion

Predictions of species coexistence constitute the building blocks for many ecological studies, such as community assembly (HilleRisLambers *et al*., 2012; Kraft *et al*., 2015; Grainger *et al*., 2019), the evolution of competitive communities (Letten *et al*., 2021; Pastore *et al*.,2021; Germain *et al*., 2022), or the role of species richness in ecosystem functioning (Godoy *et al*., 2020). Many of these studies rely on mathematical models as the basis of their predictions. Species coexistence is determined by many processes acting simultaneously, and studying it often involves a process of abstraction from ecological reality to mathematical objects such as phenomenological models (Levins, 2006). Structural sensitivity is likely to arise when these processes are summarized into equations after adopting assumptions regarding the complexity of the biological system (Aldebert & Stouffer, 2018). This makes predictions of species coexistence made with phenomenological models particularly vulnerable, especially when the model is an extreme simplification of a much more complex phenomena (Aldebert *et al*., 2018; Stouffer, 2022). Our study has shown that different phenomenological models can enhance or diminish the effect of environmental context dependency and parameter sensitivity, and thus our predictions of species coexistence. Overall, we argue that the interplay between different sources of uncertainty should not be ignored when we make model-based predictions, be they for the outcome of competition between plants or any other complex and emergent ecological phenomenon.

## Acknowledgments

We thank Trace Martyn for assistance in the field and with data collection. We would also like to thank Rogini Runghen & Hao Ran Lai for suggestions that improved the manuscript and to Hao Ran Lai for his specific input on implementing the Bayesian model fitting. ACL and DBS acknowledge the support of a Rutherford Discovery Fellowship (to DBS), administered by the Royal Society Te Apārangi; ACL, MLM, MMM, and DBS acknowledge support from the Marsden Fund Council from New Zealand Government funding, which is also managed by the Royal Society Te Apārangi (16-UOC-008 awarded to DBS & MMM); AP, MMM, and DBS thank the support provided by the Australian Research Council (DP170100837 awarded to MMM & DBS); MLM acknowledges a University of Canterbury Doctoral Scholarship. We followed the first-last-author-emphasis (FLAE) convention when determining author order (Tscharntke et al., 2007), and authorship was decided in accordance with the recommendations of the Vancouver Convention.

## References

Adler, P.B., Smull, D., Beard, K.H., Choi, R.T., Furniss, T., Kulmatiski, A., Meiners, J.M., Tredennick, A.T. & Veblen, K.E. (2018). Competition and coexistence in plant communities: intraspecific competition is stronger than interspecific competition. Ecology Letters, 21, 1319–1329.

Aldebert, C., Kooi, B., Nerini, D. & Poggiale, J. (2018). Is structural sensitivity a problem of oversimplified biological models? insights from nested dynamic energy budget models. Journal of Theoretical Biology, 448, 1–8.

Aldebert, C., Nerini, D., Gauduchon, M. & Poggiale, J. (2016). Does structural sensitivity alter complexity-stability relationships? Ecological Complexity, 28, 104–112.

Aldebert, C. & Stouffer, D.B. (2018). Community dynamics and sensitivity to model structure: towards a probabilistic view of process-based model predictions. Journal of the Royal Society Interface, 15, 20180741.

Barabás, G., D’Andrea, R. & Stump, S.M. (2018). Chesson’s coexistence theory. Ecological Monographs, 88, 277–303.

Beverton, R.J.H. & Holt, S.J. (1957). Dynamics of Exploited Fish Populations. vol. XIX of Fishery Investigations Series II. Her Majesty’s Stationery Office, London, UK.

Bimler, M.D., Stouffer, D.B., Lai, H.R. & Mayfield, M.M. (2018). Accurate predictions of coexistence in natural systems require the inclusion of facilitative interactions and environmental dependency. Journal of Ecology, 106, 1839–1852.

Bolker, B.M. (2008). Ecological Models and Data in R. Princeton University Press, Princeton, NJ, USA.

Broekman, M.J.E., Muller-Landau, H.C., Visser, M.D., Jongejans, E., Wright, S.J. & de Kroon, H. (2019). Signs of stabilisation and stable coexistence. Ecology Letters, 22, 1957–1975.

Brooker, R.W., Maestre, F.T., Callaway, R.M., Lortie, C.L., Cavieres, L.A., Kunstler, G., Liancourt, P., Tielbörger, K., Travis, J.M.J., Anthelme, F., Armas, C., Coll, L., Corcket, E., Delzon, S., Forey, E., Kikvidze, Z., Olofsson, J., Pugnaire, F., Quiroz, C.L., Saccone, P., Schiffers, K., Seifan, M., Touzard, B. & Michalet, R. (2008). Facilitation in plant communities: the past, the present, and the future. Journal of Ecology, 96, 18–34.

Bürkner, P.C. (2017). Advanced Bayesian Multilevel Modeling with the R Package brms. arXiv:1705.11123 [stat]. ArXiv: 1705.11123.

Callaway, R.M., Brooker, R.W., Choler, P., Kikvidze, Z., Lortie, C.J., Michalet, R., Paolini, L., Pugnaire, F.I., Newingham, B., Aschehoug, E.T., Armas, C., Kikodze, D. & Cook, B.J. (2002). Positive interactions among alpine plants increase with stress. Nature, 417, 844–848.

Case, T.J. (1999). An Illustrated Guide to Theoretical Ecology. Oxford University Press, Oxford, UK.

Chamberlain, S.A., Bronstein, J.L. & Rudgers, J.A. (2014). How context dependent are species interactions? Ecology Letters, 17, 881–890.

Chesson, P. (2000). General theory of competitive coexistence in spatially-varying environments. Theoretical Population Biology, 58, 211–237.

Chesson, P. (2018). Updates on mechanisms of maintenance of species diversity. Journal of Ecology, 106, 1773–1794.

Cohen, D. (1966). Optimizing reproduction in a randomly varying environment. Journal of Theoretical Biology, 12, 119–129.

Connell, J.H. (1990). Apparent versus “real” competition in plants. In: Perspectives in Plant Competition (eds. Grace, J. & Tilman, D.). Academic Press, Inc., San Diego, CA, USA, pp. 9–26.

Cordoleani, F., Nerini, D., Gauduchon, M., Morozov, A. & Poggiale, J.C. (2011). Structural sensitivity of biological models revisited. Journal of Theoretical Biology, 283, 82–91.

Craine, J.M. & Dybzinski, R. (2013). Mechanisms of plant competition for nutrients, water and light. Functional Ecology, 27, 833–840.

Dwyer, J.M., Hobbs, R.J., Wainwright, C.E. & Mayfield, M.M. (2015). Climate moderates release from nutrient limitation in natural annual plant communities. Global Ecology and Biogeography, 24, 549–561.

Dybzinski, R. & Tilman, D. (2007). Resource use patterns predict long-term outcomes of plant competition for nutrients and light. The American Naturalist, 170, 305–318.

Fussmann, G.F. & Blasius, B. (2005). Community response to enrichment is highly sensitive to model structure. Biology Letters, 1, 9–12.

García-Callejas, D., Godoy, O. & Bartomeus, I. (2020). cxr: A toolbox for modelling species coexistence in R. Methods in Ecology & Evolution, 11, 1221–1226.

Germain, R.M., Urquhart-Cronish, M., Jones, N.T., Mayfield, M.M. & Raymundo, M. (2022). The strength and direction of local (mal)adaptation depends on neighbour density and the environment. Journal of Ecology, 110, 514–525.

Godoy, O., Gómez-Aparicio, L., Matías, L., Pérez-Ramos, I.M. & Allan, E. (2020). An excess of niche differences maximizes ecosystem functioning. Nature Communications, 11, 1–10.

Godoy, O., Kraft, N.J.B. & Levine, J.M. (2014). Phylogenetic relatedness and the determinants of competitive outcomes. Ecology Letters, 17, 836–844.

Godoy, O. & Levine, J.M. (2014). Phenology effects on invasion success: insights from coupling field experiments to coexistence theory. Ecology, 95, 726–736.

Godwin, C.M., Chang, F.H. & Cardinale, B.J. (2020). An empiricist’s guide to modern coexistence theory for competitive communities. Oikos, 129, 1109–1127.

Goldberg, D.E. (1990). Components of resource competition in plant communities. In: Perspectives in Plant Competition (eds. Grace, J. & Tilman, D.). Academic Press, Inc., San Diego, CA, USA, pp. 27–49.

Grainger, T.N., Letten, A.D., Gilbert, B. & Fukami, T. (2019). Applying modern coexistence theory to priority effects. Proceedings of the National Academy of Sciences, 116, 6205–6210.

Hart, S.P., Freckleton, R.P. & Levine, J.M. (2018). How to quantify competitive ability. Journal of Ecology, 106, 1902–1909.

Hilborn, R.A.Y. & Mangel, M.A.R.C. (1997). The Ecological Detective: Confronting Models with Data. Princeton University Press, Princeton, NJ, USA.

HilleRisLambers, J., Adler, P.B., Harpole, W., Levine, J.M. & Mayfield, M.M. (2012). Rethinking community assembly through the lens of coexistence theory. Annual Review of Ecology, Evolution, and Systematics, 43, 227–248.

Houlahan, J.E., McKinney, S.T. & Rochette, R. (2015). On theory in ecology: Another perspective. BioScience, 65, 341–342.

Inouye, B.D. (2001). Response surface experimental designs for investigating interspecific competition. Ecology, 82, 2696–2706.

Jørgensen, S.E. & Bendoricchio, G. (2001). Fundamentals of Ecological Modelling. vol. 21. Elsevier.

Klir, G.J. (1985). Architecture of Systems Problem Solving. Plenum Press, New York, NY, USA.

Kraft, N.J., Adler, P.B., Godoy, O., James, E.C., Fuller, S. & Levine, J.M. (2015). Community assembly, coexistence and the environmental filtering metaphor. Functional Ecology, 29, 592–599.

Lai, H.R., Chong, K.Y., Yee, A.T.K., Mayfield, M.M. & Stouffer, D.B. (2022). Non-additive biotic interactions improve predictions of tropical tree growth and impact community size structure. Ecology, 103, e03588.

Lanuza, J.B., Bartomeus, I. & Godoy, O. (2018). Opposing effects of floral visitors and soil conditions on the determinants of competitive outcomes maintain species diversity in heterogeneous landscapes. Ecology Letters, 21, 865–874.

Law, R. & Watkinson, A.R. (1987). Response-surface analysis of two-species competition: an experiment on *Phleum arenarium* and *Vulpia fasciculata*. Journal of Ecology, 75, 871–886.

Letten, A.D., Hall, A.R. & Levine, J.M. (2021). Using ecological coexistence theory to understand antibiotic resistance and microbial competition. Nature Ecology & Evolution, 5, 431–441.

Letten, A.D., Ke, P.J. & Fukami, T. (2017). Linking modern coexistence theory and contemporary niche theory. Ecological Monographs, 87, 161–177.

Levine, J.M. & HilleRisLambers, J. (2009). The importance of niches for the maintenance of species diversity. Nature, 461, 254–257.

Levins, R. (2006). Strategies of abstraction. Biology and Philosophy, 21, 741–755.

Maestre, F.T., Callaway, R.M., Valladares, F. & Lortie, C.J. (2009). Refining the stress-gradient hypothesis for competition and facilitation in plant communities. Journal of Ecology, 97, 199–205.

Maestre, F.T., Valladares, F. & Reynolds, J.F. (2005). Is the change of plant–plant interactions with abiotic stress predictable? A meta-analysis of field results in arid environments. Journal of Ecology, 93, 748–757.

Marquet, P.A., Allen, A.P., Brown, J.H., Dunne, J.A., Enquist, B.J., Gillooly, J.F., Gowaty, P.A., Harte, J., Hubbell, S.P., Okie, J.G., Ostling, A., Ritchie, M., Storch, D. & West, G.B. (2015). On the importance of first principles in ecological theory development. BioScience, 65, 342–343.

Martyn, T.E., Stouffer, D.B., Godoy, O., Bartomeus, I., Pastore, A.I. & Mayfield, M.M. (2021). Identifying “useful” fitness models: Balancing the benefits of added complexity with realistic data requirements in models of individual plant fitness. The American Naturalist, 197, 415–433.

Matías, L., Godoy, O., Gómez-Aparicio, L. & Pérez-Ramos, I.M. (2018). An experimental extreme drought reduces the likelihood of species to coexist despite increasing intransitivity in competitive networks. Journal of Ecology, 106, 826–837.

Mayfield, M.M. & Stouffer, D.B. (2017). Higher-order interactions capture unexplained complexity in diverse communities. Nature Ecology & Evolution, 1, 0062.

McElreath, R. (2018). Statistical Rethinking: A Bayesian Course with Examples in R and Stan. Chapman and Hall/CRC.

Myerscough, M., Darwen, M. & Hogarth, W. (1996). Stability, persistence and structural stability in a classical predator-prey model. Ecological Modelling, 89, 31–42.

O’Hara, R.B. & Kotze, D.J. (2010). Do not log-transform count data. Methods in Ecology & Evolution, 1, 118–122.

Pastore, A.I., Barabás, G., Bimler, M.D., Mayfield, M.M. & Miller, T.E. (2021). The evolution of niche overlap and competitive differences. Nature Ecology & Evolution, 5, 330–337.

Pickett, S.T. (1980). Non-equilibrium coexistence of plants. Bulletin of the Torrey Botanical Club, pp. 238–248.

Poggiale, J.C., Baklouti, M., Queguiner, B. & Kooijman, S. (2010). How far details are im-portant in ecosystem modelling: the case of multi-limiting nutrients in phytoplankton–zooplankton interactions. Philosophical Transactions of the Royal Society B: Biological Sci-ences, 365, 3495–3507.

Poorter, H. & Lambers, H. (1986). Growth and competitive ability of a highly plastic and a marginally plastic genotype of Plantago major in a fluctuating environment. Physiologia Plantarum, 67, 217–222.

R Core Team (2013). R: A Language and Environment for Statistical Computing. R Foundation for Statistical Computing, Vienna, Austria.

Rao, C., Toutenberg, H., Shalabh & Heumann, C. (2010). Linear Models and Generalizations. Springer, Berlin.

Ricker, W.E. (1954a). Effects of compensatory mortality upon population abundance. The Journal of Wildlife Management, 18, 45–51.

Ricker, W.E. (1954b). Stock and recruitment. J. Fish. Res. Board Canada, 11, 559–623.

Rohr, R.P., Saavedra, S. & Bascompte, J. (2014). On the structural stability of mutualistic systems. Science, 345.

Saavedra, S., Rohr, R.P., Bascompte, J., Godoy, O., Kraft, N.J.B. & Levine, J.M. (2017). A structural approach for understanding multispecies coexistence. Ecological Monographs, 87, 470–486.

Song, C., Rohr, R.P. & Saavedra, S. (2018). A guideline to study the feasibility domain of multi-trophic and changing ecological communities. Journal of Theoretical Biology, 450, 30–36.

Song, C., Von Ahn, S., Rohr, R.P. & Saavedra, S. (2020). Towards a probabilistic under-standing about the context-dependency of species interactions. Trends in Ecology & Evolution, 35, 384–396.

Stouffer, D.B. (2022). A critical examination of models of annual-plant population dynamics and density-dependent fecundity. Methods in Ecology & Evolution, 13, 2516–2530.

Stouffer, D.B., Wainwright, C.E., Flanagan, T. & Mayfield, M.M. (2018). Cyclic population dynamics and density-dependent intransitivity as pathways to coexistence between co-occurring annual plants. Journal of Ecology, 106, 838–851.

Terry, J.C.D., Chen, J. & Lewis, O.T. (2021). Natural enemies have inconsistent impacts on the coexistence of competing species. Journal of Animal Ecology, 90, 2277–2288.

Towers, I.R., Bowler, C.H., Mayfield, M.M. & Dwyer, J.M. (2020). Requirements for the spatial storage effect are weakly evident for common species in natural annual plant assemblages. Ecology, 101, e03185.

Towers, I.R., Merritt, D.J., Erikson, T.E., Mayfield, M.M. & Dwyer, J.M. (2022). Variable seed bed microsite conditions and light influence germination in Australian winter annuals. Oecologia, 198, 865–875.

Tscharntke, T., Hochberg, M.E., Rand, T.A., Resh, V.H. & Krauss, J. (2007). Author sequence and credit for contributions in multiauthored publications. PLOS Biology, 5, e18.

Van Dyke, M.N., Levine, J.M. & Kraft, N.J.B. (2022). Small rainfall changes drive substantial changes in plant coexistence. Nature, 611, 507–511.

Vehtari, A., Gelman, A. & Gabry, J. (2017). Practical Bayesian model evaluation using leave-one-out cross-validation and WAIC. Statistics and Computing, 27, 1413–1432.

Vehtari, A., Gelman, A., Simpson, D., Carpenter, B. & Bürkner, P.C. (2021). Rank-normalization, folding, and localization: An improved Rhat for assessing convergence of mcmc. Bayesian Analysis, 16, 667–718.

Villarreal-Barajas, T. & Martorell, C. (2009). Species-specific disturbance tolerance, competition and positive interactions along an anthropogenic disturbance gradient. Journal of Vegetation Science, 20, 1027–1040.

Wainwright, C.E., Dwyer, J.M., Hobbs, R.J. & Mayfield, M.M. (2017a). Diverse outcomes of species interactions in an invaded annual plant community. Journal of Plant Ecology, 10, 918–926.

Wainwright, C.E., Dwyer, J.M. & Mayfield, M.M. (2017b). Effects of exotic annual grass litter and local environmental gradients on annual plant community structure. Biological Invasions, 19, 479–491.

Wainwright, C.E., HilleRisLambers, J., Lai, H.R., Loy, X. & Mayfield, M.M. (2019). Distinct responses of niche and fitness differences to water availability underlie variable coexistence outcomes in semi-arid annual plant communities. Journal of Ecology, 107, 293–306.

Watkinson, A.R. (1980). Density-dependence in single-species populations of plants. Journal of Theoretical Biology, 83, 345–357.

Wood, S.N. & Thomas, M.B. (1999). Super-sensitivity to structure in biological models. Proceedings of the Royal Society of London. Series B: Biological Sciences, 266, 565–570.

Zeigler, B.P., Praehofer, H. & Kim, T. (2000). Theory of Modeling and Simulation. Academic Press, San Diego, CA, USA.

Zepeda, V. & Martorell, C. (2019). Fluctuation-independent niche differentiation and relative non-linearity drive coexistence in a species-rich grassland. Ecology, 100, e02726.

