## Supporting Information for "Environmental context, parameter sensitivity and structural sensitivity impact predictions of annual-plant coexistence"

Cervantes-Loreto et al.

#### Appendix S1 The biologically constrained feasibility domain, $\beta$

We used a Monte-Carlo integration approach to estimate the size of area corresponding to the biologically constrained feasibility domain ($\beta$); that is, the space of vital rates that allow both species to have positive equilibrium abundances given the interaction coefficients, abundance constraints, and model-based constraints. The objective of our approach is to generate a large set of randomly sampled vital rates $(r_i, r_j)$ to represent values in the biologically constrained feasibility domain.

With two species, any potential pair of vital rates can be thought of points falling on a circle of radius $R = \sqrt{r_i^2 + r_j^2}$. As described in the main text, larger and larger radii $R$ generally correspond to larger and larger equilibrium densities $N_i^*$ and $N_j^*$ predicted by the population-dynamics models. Since the feasibility of a vector $\tilde{r}$ guarantees feasibility of every proportional vector $x\tilde{r}$ (where $x$ is any scalar value), in the most common implementation the value of $R$ is fixed to 1; this converts the two-dimensional domain in $(r_i, r_j)$ space to an angle over which the vital rates may vary (Rohr *et al.*, 2014; Saavedra *et al.*, 2017). Otherwise, the domain will mathematically have an infinitely large area. In contrast, our approach imposes an upper bound on the densities that are regarded as biologically and empirically sensible. This implies that there will always be a maximum radius $R_{\max}$ beyond which even feasible equilibria will no longer respect the constraints we impose on equilibrium densities. Unfortunately, there is no guarantee that the arbitrary, albeit convenient, selection of $R = 1$ will always satisfy these constraints.

On the other hand, every biologically sensible feasible equilibrium corresponds to a unique set of species vital rates. This implies that we can uniformly sample feasible equilibria and from these determine the space of vital rates that is consistent with them. We therefore developed an approach fashioned after Monte-Carlo integration that proceeds as follows. We first generate a random equilibrium point $(n_i^*, n_j^*)$, where $n_i^* = g_i N_i^*$ and $n_j^* = g_j N_j^*$ are the equilibrium densities of plants. The values of $n_i^*$ and $n_j^*$ are sampled uniformly in the ranges $[0, n_{i,\max}]$ and $[0, n_{j,\max}]$, respectively. For all of the models studied here, the vector of species vital rates can be expressed as a linear function of species equilibrium densities such that the two equations:

$$\begin{aligned} r_i &= \alpha_{ii} n_i^* + \alpha_{ij} n_j^* \\ r_j &= \alpha_{ji} n_i^* + \alpha_{jj} n_j^* \end{aligned} \tag{S1}$$

provide a mapping from $(n_i^*, n_j^*)$ to $(r_i, r_j)$. Of course, not every feasible equilibria is necessarily biological because of the model-based constraints. We therefore only retained the paired vital rates $(r_i, r_j)$ if they also fell within the model-based constraints on $r_i$ and $r_j$ for the two species (Table S1). This method allowed us to sample points that were feasible, biologically plausible, and were consistent with the model constraints. We set our integration routine to sample random points until we had at least 50000 points inside the feasible and biologically plausible space. All calculations were done in the programming language $R$ (R Core Team, 2013).

Once we had a sample of feasible and biologically constrained vital rates ($\beta$), we determined

**Table S1:** Equilibrium vital rates and model-based constraints thereof for the two models of density-dependent fecundity we used used to make coexistence predictions. Vital rates are given by solving each model's equilibrium conditions. Upper and lower bounds are the result of how bounds on $g_i$, $s_i$, and $\lambda_i$ come together to impact the values vital rate can take while maintaining biological interpretability.

| Model | Vital rate, $r_i$ | Lower bound | Upper bound |
|---|---|---|---|
| Beverton–Holt | $-1 + \left(\frac{g_i \lambda_i}{1-(1-g_i)s_i}\right)$ | $-1$ | $\infty$ |
| Ricker | $\ln\left(\frac{g_i \lambda i}{1-(1-g_i)s_i}\right)$ | $-\infty$ | $\infty$ |

the points that comprise the convex hull around them using the function **chull** from the package **grDevices**. We then calculated the size of the area described by the convex hull, using the function **Polygon** from the package **sp**. To test the assumption that the feasible and biologically constrained points are accurately described by a convex hull, we tested whether any discarded points fell inside the area described by the vertices using the function **point.in.polygon** from the package **sp**. None of our sampled feasibility domains violated this test and hence we concluded that treating the biologically constrained feasibility domain as a convex hull was appropriate.

#### Appendix S2 The area in monoculture, $\gamma$

To determine the parameter space where both species can grow in monoculture given abundance and model-based constraints, we first determined the extremal values of species vital rates $r_{i,\max}$ and $r_{j,\max}$ that correspond to $r_{i,\max} = \alpha_{ii} n_{i,\max}$ and $r_{j,\max} = \alpha_{jj} n_{j,\max}$, respectively. Note that the values of species vital rates $r_{i,\max}$ and $r_{j,\max}$ can take negative values if intraspecific competition coefficients are negative (which would imply facilitative interactions). Without model-based constraints, the range of vital-rate parameter space is a rectangle with the coordinate $(r_i = 0, r_j = 0)$ on one corner and these extremal values on the other. However, the inclusion of model-based constraints implies that the biological feasible and model-consistent region of vital rates is given by

$$r_i \in \begin{cases} [\max\{r_{i,\max}, r_{i,\text{lower}}\}, 0], & \text{if } r_{i,\max} < 0 \\ [0, \min\{r_{i,\max}, r_{i,\text{upper}}\}], & \text{otherwise} \end{cases} \tag{S2}$$

where $r_{i,\text{lower}}$ and $r_{i,\text{upper}}$ are the lower and upper limits of species vital rates given the models used to quantify density dependence for each species (Table 1 in the main text). An equivalent expression for species $j$ can be obtained by swapping all $i$ subscripts for $j$ subscripts. Given the bounds on $r_i$ and $r_j$, we then calculate the area $\gamma$ of the rectangle as the product of the length of both sides.

### Appendix S3 Germination rate, seed survival rate, and maximum plant abundances

We relied on independent empirical measures of seed germination rate, seed survival rate, and maximum expected plant abundances for both *Goodenia rosea* and *Trachymene cyanopetala*. We obtained values of seed survival rate and germination rate from a previous study that looked at species-level metrics in our study system (Towers *et al.*, 2022). We determined the maximum abundance of seeds based on empirical observations of individual plants of both species within a 7.5 cm radius. Both *G. rosea* and *T. cyanopetala* rarely exceed abundances larger than 100 plant individuals within a 7.5 cm radius (Trace E. Martyn, personal communication). Nevertheless, we decided to use a threshold that was five times larger than these empirical observations so that the abundance constraints would be as conservative as possible. The resulting seed survival rates, germination rates, and maximum abundances we used across our analyses were:

| Species | Germination rate, $g$ | Seed survival rate, $s$ | Maximum plant abundance, $n_{\max}$ | Maximum seed abundance, $N_{\max}$ |
|---|---|---|---|---|
| *Goodenia rosea* | 0.964 | 0.965 | 500 | 518 |
| *Trachymene cyanopetala* | 0.474 | 0.969 | 500 | 1054 |

### Appendix S4 Posterior distributions of model parameters

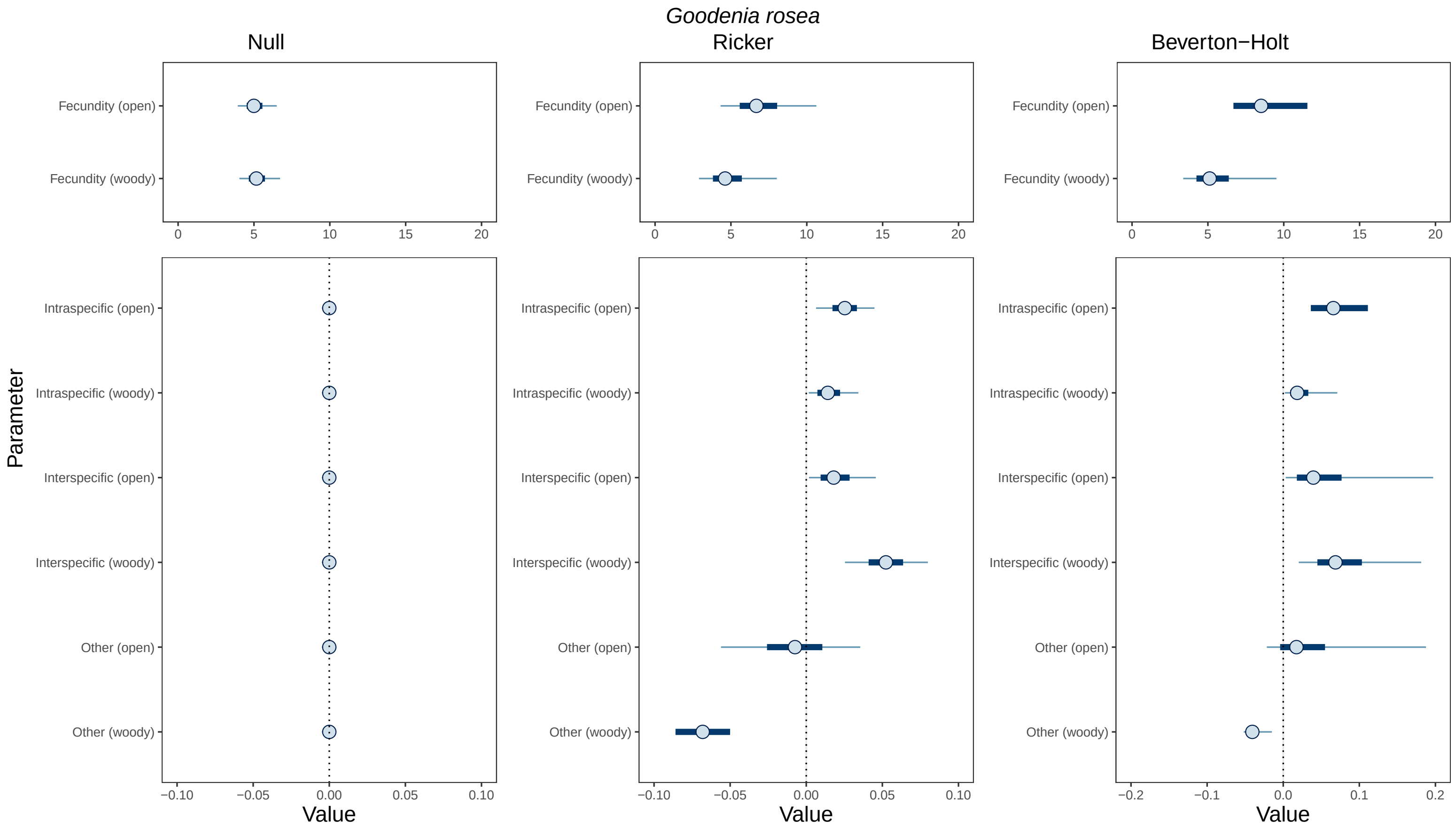


**Figure S1:** The posterior distributions of competition coefficients and intrinsic fecundity inferred when using the Null, Ricker, and Beverton–Holt models to describe density-dependent plant fecundity for *Goodenia rosea*. Blue circles indicate parameters' median estimates, dark blue bars correspond to the inner 50% probability mass of each parameter's posterior distribution, and thin lines extend out to their 90% interval. Finally, we denote the environment where interactions took place to estimate parameter values with *open* and *woody*, as we do in the main text.

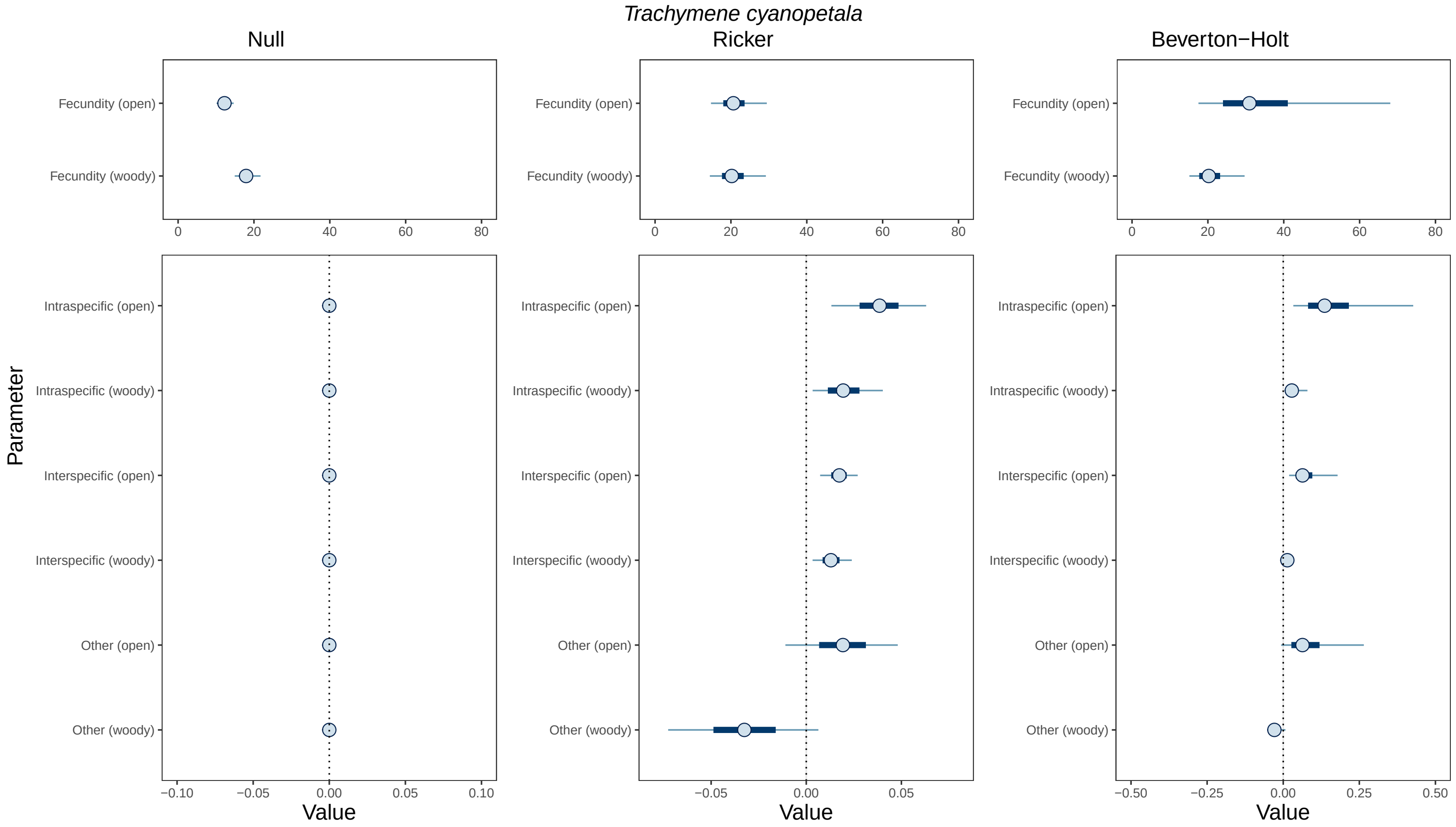


**Figure S2:** The posterior distributions of competition coefficients and intrinsic fecundity inferred when using the Null, Ricker, and Beverton–Holt model to describe density-dependent plant fecundity for *Trachymene cyanopetala*. Blue circles indicate parameters' median estimates, dark blue bars correspond to the inner 50% probability mass of each parameter's posterior distribution, and thin lines extend out to their 90% interval. Finally, we denote the environment where interactions took place to estimate parameter values with *open* and *woody*, as we do in the main text.

#### Appendix S5 Posterior distributions of feasibility domains

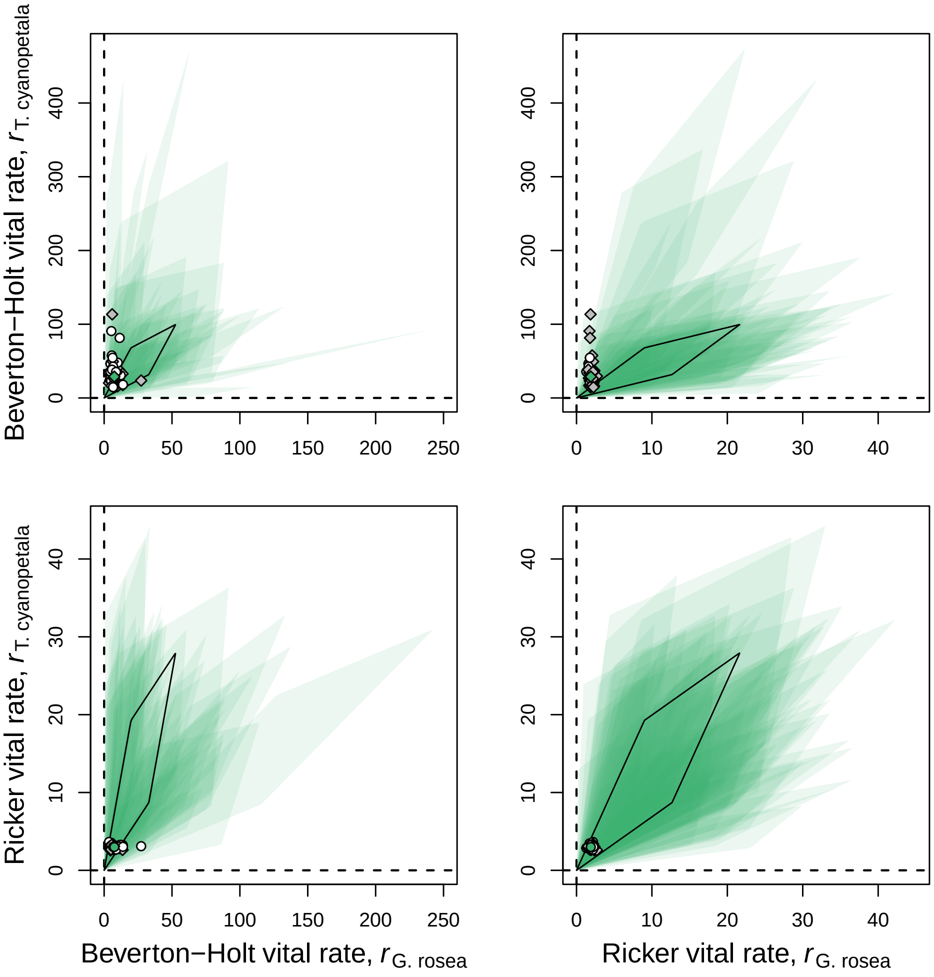


**Figure S1:** Posterior distributions of the biologically constrained feasibility domains and species' inferred vital rates for different model combinations in the *open* environment. For a representative example of 50 posterior draws, we plot both the biologically constrained feasibility domain and corresponding vital rate vector. Each feasibility domain is shaded in light green with transparency, such that areas where multiple domains overlap are indicated with darker shading. Since it is difficult to associate each pair of vital rates with each feasibility domain across 50 separate draws, white circles indicate a set of vital rates that fell inside the feasibility domain (i.e., predict coexistence) and gray circles indicate a set of vital rates that fell outside the feasibility domain (i.e., predict competitive exclusion). The solid black line also indicates the outline of the biologically constrained feasibility domain using median parameter values. The green filled point indicates the median inferred vital rates, and the shape of that point indicates the predicted outcome using median parameter values (circle = coexistence; diamond = competitive exclusion).

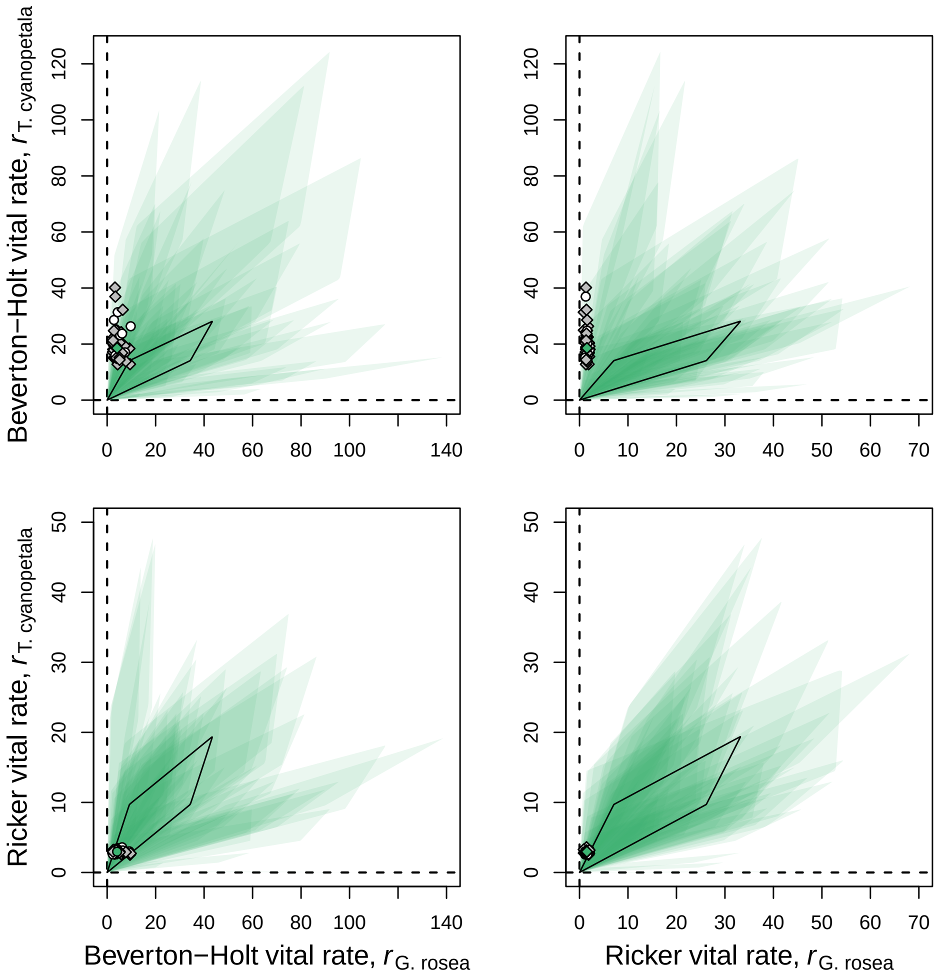


**Figure S2:** Posterior distributions of the biologically constrained feasibility domains and species' inferred vital rates for different model combinations in the *woody* environment. For a representative example of 50 posterior draws, we plot both the biologically constrained feasibility domain and corresponding vital rate vector. Each feasibility domain is shaded in light green with transparency, such that areas where multiple domains overlap are indicated with darker shading. Since it is difficult to associate each pair of vital rates with each feasibility domain across 50 separate draws, white circles indicate a set of vital rates that fell inside the feasibility domain (i.e., predict coexistence) and gray circles indicate a set of vital rates that fell outside the feasibility domain (i.e., predict competitive exclusion). The solid black line also indicates the outline of the biologically constrained feasibility domain using median parameter values. The green filled point indicates the median inferred vital rates, and the shape of that point indicates the predicted outcome using median parameter values (circle = coexistence; diamond = competitive exclusion).

#### Supplementary References

R Core Team (2013). *R: A Language and Environment for Statistical Computing*. R Foundation for Statistical Computing, Vienna, Austria.

Rohr, R.P., Saavedra, S. & Bascompte, J. (2014). On the structural stability of mutualistic systems. *Science*, 345.

Saavedra, S., Rohr, R.P., Bascompte, J., Godoy, O., Kraft, N.J.B. & Levine, J.M. (2017). A structural approach for understanding multispecies coexistence. *Ecological Monographs*, 87, 470–486.

Towers, I.R., Merritt, D.J., Erikson, T.E., Mayfield, M.M. & Dwyer, J.M. (2022). Variable seed bed microsite conditions and light influence germination in Australian winter annuals. *Oecologia*, 198, 865–875.
